# A genetic anemia disentangles the dual effect of chronically reduced thrombopoietin on hematopoietic stem cells and bone marrow niche

**DOI:** 10.64898/2026.10.08.757539

**Authors:** Annamaria Aprile, Alessandro Malara, Mariangela Storto, Alessandro Gulino, Valeria Cancila, Gaia Morello, Silvia Sighinolfi, Laura Raggi, Stefano Beretta, Ivan Merelli, Sarah Marktel, Maurilio Ponzoni, Claudio Tripodo, Alessandra Balduini, Giuliana Ferrari

## Abstract

Many studies unraveled the reciprocal regulation between hematopoietic stem cells (HSC) and the bone marrow (BM) niche by using transgenic knock-out or reporter mice, often overlooking the pathophysiological relevance and the existence of compensatory mechanisms. Here, we exploit the *th3* murine model of a genetic anemia, beta-thalassemia (BThal), to investigate the dual impact of chronically reduced thrombopoietin (TPO) production on HSC and megakaryocytic BM niche. Decreased systemic TPO downregulates stemness programs in HSC and impairs their commitment towards megakaryocytes (Mk). We uncover dysmegakaryopoiesis in the BM from BThal patients and mice with defective Mk maturation and reduced production of niche factors, altering the HSC-niche crosstalk. Notably, *in vivo* stimulation of the TPO signaling restores HSC quiescence, Mk maturation and niche support. These findings demonstrate that TPO regulates HSC through both direct and niche-mediated routes, underscoring the value of using a genetic disease model to disentangle general molecular mechanisms.

## INTRODUCTION

Under homeostatic conditions, the adult hematopoietic system is maintained by the hematopoietic stem cells (HSC), which lie at the apex of the hierarchical process of hematopoiesis and reside within the bone marrow (BM) in a specialized environment, termed niche^1^. HSC are highly quiescent, only occasionally enter cell cycle and throughout an individual’s lifetime, they self-renew and differentiate to produce multilineage blood cells. HSC behavior is governed by interactions with the complex niche microenvironment, providing growth factors, cytokines and other stimuli to modulate HSC fate. Several cellular components of the niche have been identified, including non-hematopoietic BM stromal cells, such as osteolineage, endothelial and mesenchymal stromal cells, and also differentiated hematopoietic cells, such as mature megakaryocytes (Mk), BM macrophages (Mϕ) and activated regulatory T cells^2^.

In the last decades several studies unraveled the complex regulation of HSC and the BM niche. Knocking-out or forcing the overexpression of a specific gene, indeed, helped to identify its specific role on HSC function and BM niche supporting capacity^3–6^, but risk to constrain the knowledge of cellular and molecular mechanisms in physiological and pathological conditions since in “real-world” conditions compensating molecules and pathways exist. In this context, the analysis of HSC and BM niche has been almost completely restricted to steady-state and malignancies and only recently the field of study has been extended to models of aging or acute stress stimuli, such as inflammation^7^. However, the dissection of HSC-niche crosstalk in chronic stress conditions and inherited disorders remains underexplored. The study of HSC and BM niche in these settings might be clinically relevant for transplantation and gene therapy application in hematological genetic diseases, but also could help to unmask general mechanisms, as recently demonstrated in a rare genetic disorder due to MECOM haploinsufficiency^8^.

Here, we took advantage of a model of genetic anemia, beta-thalassemia (BThal), caused by significantly reduced or absent synthesis of hemoglobin beta-chains in erythroid cells^9^. Although the genetic mutations have no direct effect on HSC, we recently demonstrated an impaired HSC function due to secondary alterations leading to a defective BM stromal niche^10–15^. In the murine model of the disease, the *Hbb^th^*^3^*^/+^*(*th3*) mouse^16^, in addition to the BM stroma, we found altered levels of multiple local and systemic factors, including reduction of systemic thrombopoietin (TPO).

TPO and its receptor, myeloid proliferative leukemia (MPL), were initially identified as the key signaling that stimulates megakaryopoiesis and platelet (Plt) production^17^. Serum TPO fluctuates in response to circulating Plt number. Unlike other cytokine-receptor pathways that trigger a single lineage of hematopoiesis, TPO-MPL signaling affects both HSC function and megakaryopoiesis. The role of TPO on HSC has been studied in constitutive and conditional knock-out models and has been demonstrated to be involved in the regulation of HSC number and quiescence^18–20^ and to modulate metabolic changes in HSC maintaining the stem cell potential^21^. In humans, patients with congenital amegakaryocytic thrombocytopenia due to loss-of-function mutations in MPL suffer a progressive decline of HSC number and subsequent BM failure^22^.

TPO regulates megakaryopoiesis and Plt production and mature Mk secrete TPO, which was thought to stimulate by a paracrine effect HSC^23^. However, recent evidence demonstrates that TPO expressed by hepatocytes and not by BM cells is responsible for HSC fate^24^. On the other hand, mature Mk regulate HSC quiescence by secreting specific cytokines, such as platelet factor 4 (PF4 or CXCL4), FGF1 and TGF-beta 1^6,25^ and in condition of defective maturation, negatively affect stem cell function^26^. Very recently the heterogeneity of Mk was dissected in terms of transcriptional profile and differentiation origin identifying Plt-producing, immune and niche-supporting Mk^27,28^.

Despite several studies unraveled the TPO function in hematopoiesis, limitations in the experimental systems have constrained our knowledge of the dual effect of chronically altered TPO on HSC and BM niche in the same setting. The study of the action of this cytokine specifically on the BM niche is lacking, as well as the simultaneous analysis of HSC in a condition of low TPO.

Here, we demonstrated that chronically reduced levels of systemic TPO impair HSC function by both a direct mechanism and an indirect one mediated by the niche-supporting Mk in a model of genetic anemia. We took advantage of the novel concept of exploiting a genetic disorder as a natural experiment to shed light into general molecular mechanisms of stem cell biology.

## RESULTS

### Reduced TPO is caused by the increased half-life of circulating Plt in th3 mice

We have recently investigated HSC function and BM microenvironment in the murine model of the genetic anemia BThal (*th3* strain) and we demonstrated reduced HSC content, quiescence and repopulating capacity as compared to wild type (wt) littermates^10^. In addition to the BM stroma, we found altered levels of multiple local and systemic factors, including an unexpected reduction of systemic TPO **(Fig. 1A)**. The halved levels of TPO in *th3* mice were due to a reduced expression by hepatocytes **(Fig. 1B)**, as liver is the main site of TPO production^24^.

**Figure 1.**
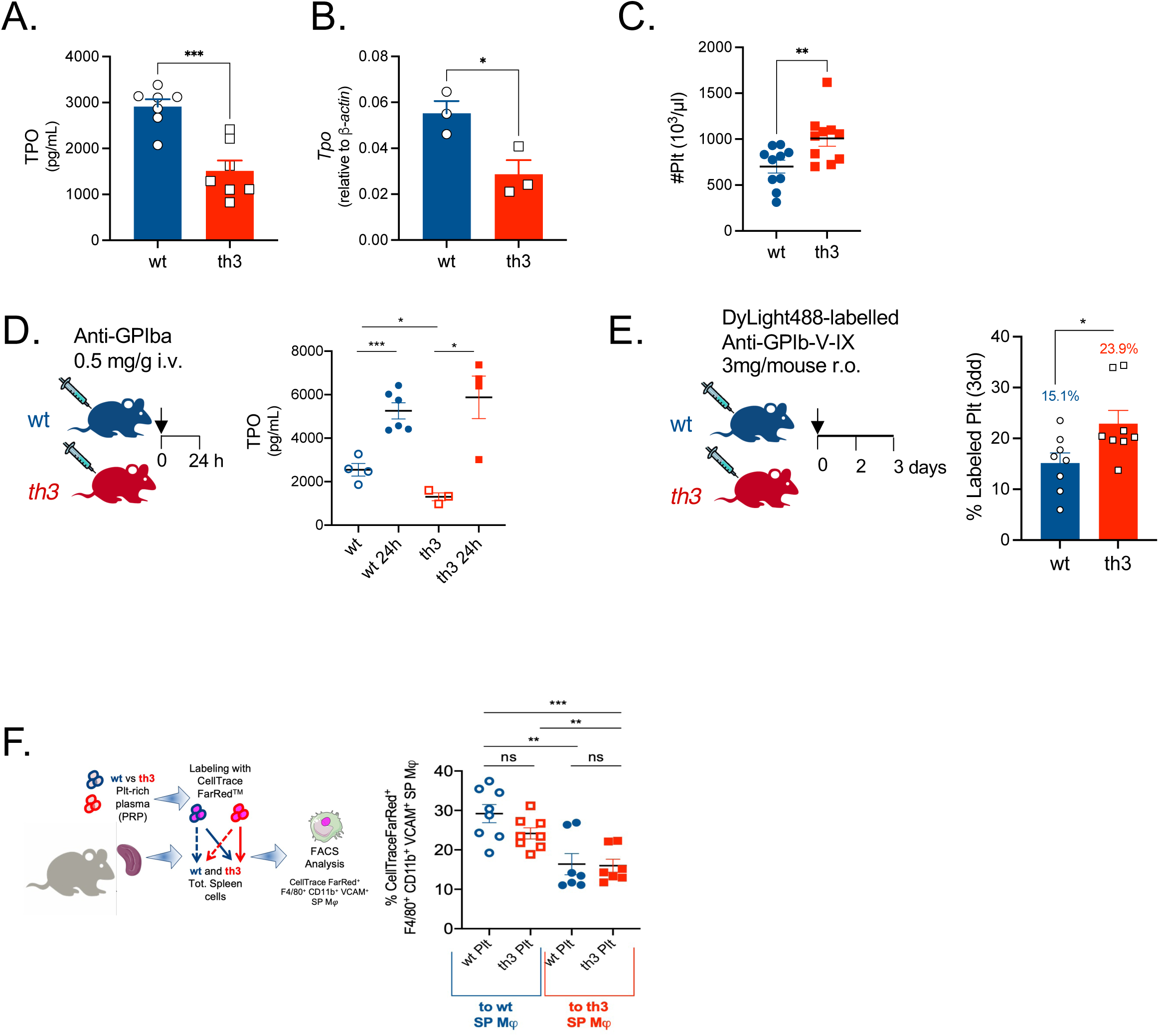
Chronically reduced TPO is caused by the increased half-life of circulating Plts in a model of genetic anemia. **A**. Serum level of TPO in *th3* and control mice (wt *n* = 7; *th3 n* = 7). **B.** Expression of *Tpo* by wt and *th3* liver cells. Fold expression to the beta-actin reference gene is reported (wt *n* = 3; *th3 n* = 3). **C.** Circulating Plt count in wt and *th3* mice by hemocytometer analysis (wt *n* = 10; *th3 n* = 10). **D.** Serum levels of TPO in wt and *th3* mice at steady-state and at 24h after acute *in vivo* Plt depletion by anti-GPIa antibody (wt *n* = 4; wt 24h *n* = 6; *th3 n* = 3; *th3* 24h *n* = 4). **E.** Frequency of labeled Plt 3 days upon injection of DyLight488-labelled anti-GPIb-V-IX antibody in wt and *th3* mice (wt *n* = 8; *th3 n* = 8). **F.** Experimental plan and flow cytometry analysis of *in vitro* Plt phagocytosis by spleen Mφ. Evaluation of the frequency of CellTrace FarRed^+^ F4/80^+^ CD11b^+^ VCAM^+^ cells from wt and *th3* spleens upon incubation with matched/mismatched labeled Plt (wt *n* = 8; *th3 n* = 7).

We explored the origin of TPO defect: since TPO fluctuates in response to Plt number, we hypothesized that TPO is downregulated in response to an increased number of circulating Plt, as confirmed by our analysis in *th3* mice **(Fig. 1C)** and reported by others also in BThal patients^29,30^. To demonstrate this feedback regulation and exclude any intrinsic defect in TPO production, we tested the levels of systemic TPO upon acute Plt depletion. Plt were depleted in *th3* mice and wt controls by a single injection of anti-GPIba antibodies and analyzed after 24h. Despite the baseline differences in TPO, *th3* mice are able to upregulate the production of this cytokine at the same level as wt animals **(Fig. 1D)**, thus excluding any intrinsic defect. We hypothesize that the Plt accumulation we described might be due to a higher half-life of *th3* Plt. Thus, we performed *in vivo* Plt labeling assays by DyLight488-labelled anti-GPIb-V-IX antibody and the labeled Plt were analyzed on whole blood stained with anti-CD61 antibody at 3 days upon injection. Consistently with our hypothesis, the results showed a higher percentage of labelling in *th3*, indicating an increased Plt half-life **(Fig. 1E)**.

The increased Plt half-life might be due to intrinsic properties of *th3* Plt or to a reduced clearance by Mϕ. In order to test the two different scenarios, we studied Plt activation and the expression of surface molecules, which make Plt more susceptible to phagocytosis by Mϕ. We evaluated by flow cytometry analysis the expression of P-selectin (CD62P) at steady-state and upon activation by *in vitro* treatment with thrombin and we found no differences between wt and *th3* samples **(Suppl. Fig. 1A)**. Moreover, levels of phosphatidylserine (Annexin V) and the “don’t eat me signal” molecule CD47 revealed that the expression of activation markers is unaltered in *th3* Plt **(Suppl. Fig. 1B,C)**, thus suggesting the lack of intrinsic Plt defects. On the other hand, as direct demonstration of altered Plt phagocytosis in *th3* animals, we performed *in vitro* phagocytosis assay by incubating Plt labelled with CellTrace FarRed^TM^ with cells from spleen, a site of Plt clearance. Matched (wt Plt + wt splenocytes; *th3* Plt + *th3* splenocytes) and mismatched (wt Plt + *th3* splenocytes; *th3* Plt + wt splenocytes) settings were evaluated and the frequency of splenic Mϕ that had phagocytosed Plt was assessed by analyzing CellTrace FarRed^+^ F4/80^+^ CD11b^+^ VCAM^+^ cells by flow cytometry. Interestingly, this assay revealed a reduced Plt phagocytosis by *th3* spleen Mϕ irrespectively from the origin of Plt **(Fig. 1F)**, further excluding any intrinsic Plt defect. The increased iron content in *th3* splenic Mϕ associated to BThal **(Suppl. Fig. 1D)** might contribute to their phagocytic defect.

### Reduced TPO response in th3 HSC

Since we have recently demonstrated that BThal HSC are more cycling and show low engraftment and repopulating capacity^10^, we investigated whether reduced TPO might contribute to impaired HSC function. Despite a normal expression of MPL receptor **(Suppl. Fig. 2A)**, analysis of RNA-seq datasets obtained from sorted Lineage^-^ Sca-1^+^ c-Kit^+^ CD48^-^ CD150^+^ HSC^10^ revealed that in *th3* HSC the TPO signaling pathway is downregulated **(Suppl. Fig. 2B)** as well as the expression of TPO target genes associated to HSC quiescence and stemness, including *Cdkn1a*, *Hoxb4* and *Hoxa9* **(Fig. 2A)**. These data are consistent with reported results of higher cycling activity of HSC in the presence of low TPO levels^19,24^ and correlate with our finding of more cycling HSC in *th3* mice^10^.

**Figure 2.**
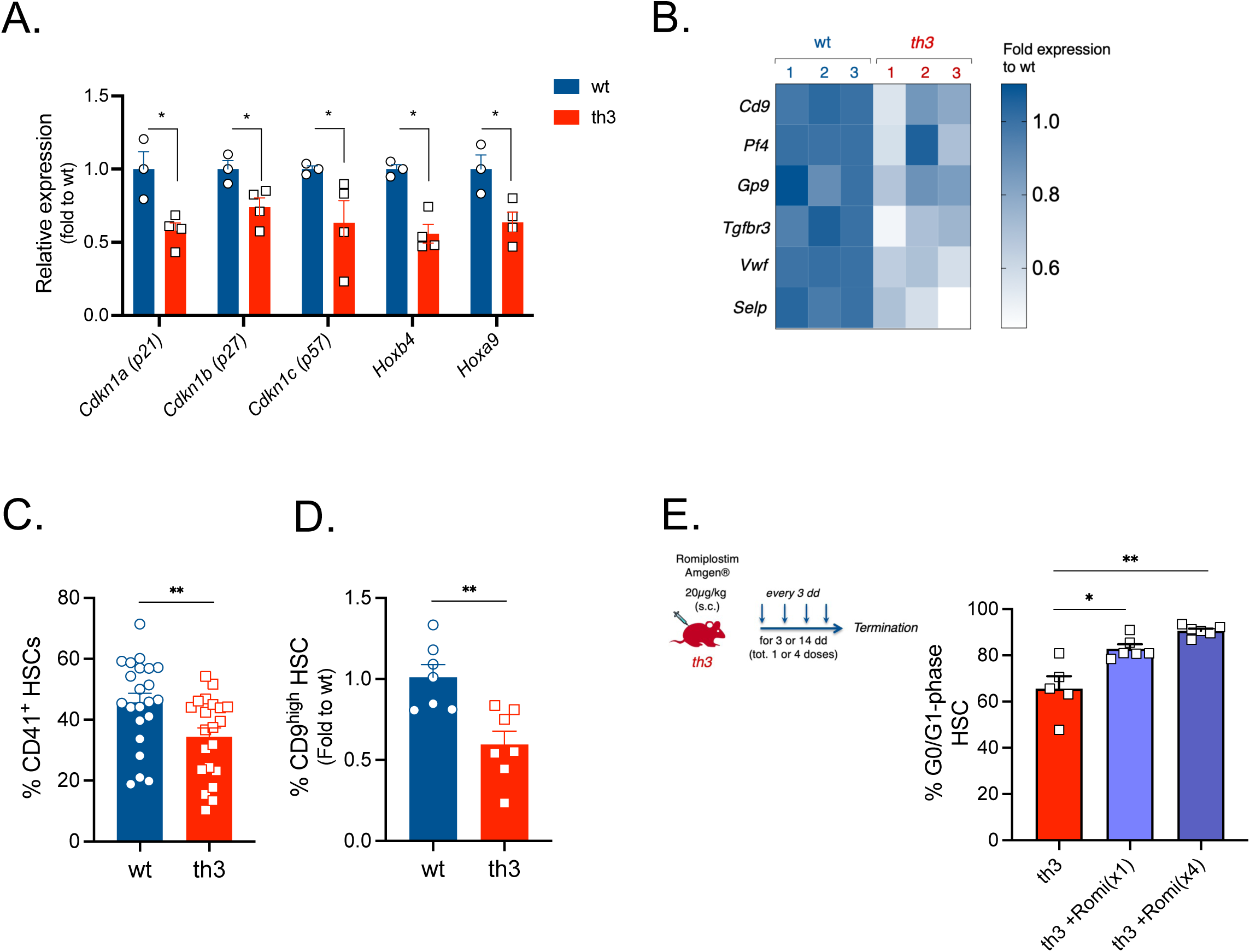
Reduced TPO response in BThal HSC. **A.** Expression levels of quiescence and stemness genes targets of TPO signaling in wt and *th3* HSC by RNA-seq analysis. Data are reported as normalized counts and expressed as fold to wt controls (wt *n* = 3; *th3 n* = 4). **B.** Gene expression heatmap of Mk-biased genes in sorted wt and *th3* HSC by RNA-seq analysis. Data are reported as fold to wt controls (wt *n* = 3; *th3 n* = 3). **C,D.** Frequency of Mk-biased CD41^+^ HSC and CD9^high^ HSC in wt and *th3* mice (wt *n* = 22 and 8; *th3 n* = 21 and 8, respectively). **E.** Frequency of quiescent HSC in G0/G1-phase of the cell cycle upon *in vivo* stimulation of TPO signaling in *th3* mice by one or 4 doses of Romiplostim (*th3 n* = 5; *th3*+Romi(x1) *n* = 6; *th3*+Romi(x4) *n* = 6).

To investigate any skewing of the Mk commitment in the more primitive hematopoietic populations we analyzed the expression of Mk and Plt genes by *th3* HSC and we found downregulated expression of genes as *Pf4*, *Gp9* and *Vwf* **(Fig. 2B)**, thus suggesting a reduced frequency of Mk-biased HSC. Consistently, immunophenotypic analysis of Mk-biased HSC expressing CD41 surface marker^31^ showed a reduced frequency of CD41^+^ HSC in *th3* mice **(Fig. 2C)**, as compared to controls. We consolidated these results by assessing a lower frequency also of the CD9^high^ Mk-biased HSC^21^ **(Fig. 2D)**.

Importantly, *in vivo* stimulation of TPO axis by Romiplostim TPO-receptor agonist (TPO-ra) in *th3* mice stimulates Plt production **(Suppl. Fig. 3A)** and restore the defective frequency and quiescence of HSC **(Suppl. Fig. 3B** and **Fig. 2E)**, thus demonstrating the contribution of defective TPO signaling in impairing HSC.

### Defective Mk maturation in th3 BM

Since TPO is also a key regulator of Mk maturation, we considered whether the reduced systemic levels in *th3* mice could affect megakaryopoiesis. *Th3* Mk showed reduced activation of TPO signaling **(Suppl. Fig. 4A)** and downregulation of transcription factors associated to megakaryopoiesis and genes related to endomitosis **(Suppl. Fig. 4B)**.

Histopathological analyses of the BM hematopoietic parenchyma of *th3* mice revealed a defect in Mk maturation, characterized by an increased frequency of morphologically immature Mk (< 10 µm diameter) and a reduced proportion of terminally mature Mk (> 25 µm diameter) (**Fig. 3A-C**). Despite the frequency and absolute counts of BM Mk detected as Lin^-^ CD41^+^ CD61^+^ cells was normal (**Suppl. Fig. 5A,B**), the maturation defects of Mk subpopulations were confirmed by the analysis of their ploidy status. Nuclear content evaluation showed a reduced frequency of mature polyploid Mk in *th3* mice and a shift towards more immature Mk, with an almost complete loss of the normal 32N profile (**Fig. 3D**). As consequence of the impaired megakaryopoiesis, we found reduced *in vivo* Plt biogenesis, assessed by daily monitoring of Plt recovery upon depletion with anti-GPIba antibody (**Suppl. Fig. 6**). Notably, *in vivo* stimulation with one or four doses of the TPO-ra Romiplostim normalized Mk diameters in *th3* mice, demonstrating that the defective Mk maturation results from impaired TPO signaling **(Fig. 3E)**.

**Figure 3.**
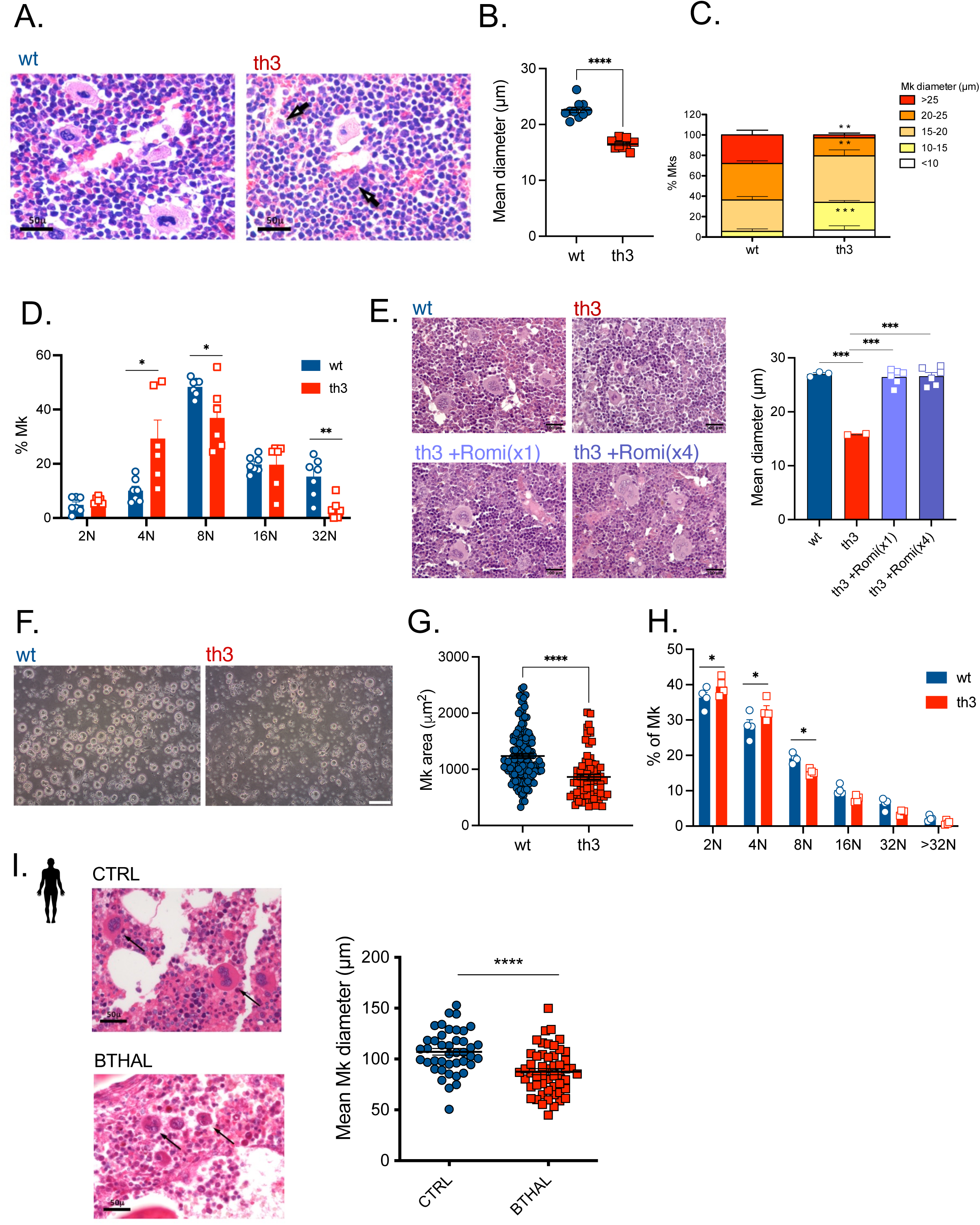
Defective Mk maturation in BThal BM. **A.** Histological analysis of BM sections (H&E). Arrows indicate morphologically immature Mk in *th3* BM. **B.** Mean diameter (µm) of wt and *th3* Mk is reported (wt *n* = 10; *th3 n* = 10). **C.** Distribution of Mk by considering their diameter (µm) is reported. Mk have been divided into 5 arbitrary classes from immature to more mature ones (<10,10-15,15-20,20-25 and >25µm diameter) (wt *n* = 5; *th3 n* = 5). **D.** Mk frequency within the total Lin^-^ CD41^+^ Mk population, based on 2N, 4N, 8N, 16N and 32N ploidy by DRAQ5^TM^ staining and cytofluorimetric analysis (wt *n* = 7; *th3 n* = 6). **E.** Representative pictures and mean diameter (µm) of Mk is reported in wt, *th3* controls and *th3* mice upon *in vivo* TPO stimulation by one or 4 doses of Romiplostim (wt *n* = 3; *th3 n* = 2; *th3*+Romi(x1) *n* = 6; *th3*+Romi(x4) *n* = 6). **F.** Representative pictures of Mk cultures from Lin^-^ cells of wt and *th3* mice upon 4dd of differentiation in the presence of TPO. **G.** Area of differentiated Mk from wt and *th3* mice (wt Mk *n* = 139; *th3* Mk *n* = 72). **H.** Ploidy levels of differentiated Mk populations by propidium Iodide staining and cytofluorimetric analysis (wt *n* = 4; *th3 n* = 4). **I.** Histological analysis of BM biopsies (H&E) from THAL and controls (CTRL). Arrows indicate morphologically immature Mk in THAL BM. Mean diameter of Mk is reported (CTRL Mk *n* = 42; BTHAL Mk *n* = 52).

We further provided evidence that *in vitro* differentiation of BM lineage-depleted hematopoietic progenitors for 4 days in the presence of recombinant TPO recapitulated the defective Mk maturation with accumulation of Mk with lower diameter and reduced ploidy **(Fig. 3F-H)**.

To further dissect *th3* megakaryopoiesis, we evaluated BM Mk progenitors, namely MkP and PreMegE cells by using the antibodies against Sca-1, c-Kit, CD150, CD16/32 and CD105 surface markers and the gating strategy reported by^32^. MkP and PreMegE cell frequencies in *th3* mice were similar to wt ones **(Suppl. Fig. 7A-C)**, as also assessed at functional level by the similar number of colonies obtained by *in vitro* colony forming unit (CFU)-Mk assay upon seeding of BM lineage-depleted cells for 6 days **(Suppl. Fig. 7D)**. Of note, analysis on BM sections derived from BThal patients confirmed defective megakaryopoiesis (**Fig. 3I**).

### Altered gene expression profile and defective BM niche support by th3 Mk

Consistently with the defective maturation of *th3* Mk, RNA-seq analysis of sorted Lineage^-^ CD41^+^ CD61^+^ Mk showed a different expression profile compared to wt. Principal component analysis (PCA) performed using the 100 most significant differentially expressed genes (DEG) highlighted the clusterization of *th3* Mk samples apart from wt controls with a variance of 53% in PC1 **(Fig. 4A)**. We identified 101 DEG (FDR <0.01) by using DESeq package with 42 upregulated and 59 downregulated genes in *th3* Mk compared to controls **(Fig. 4B** and **Suppl. Table 1)**. Gene set enrichment analysis (GSEA) employing the biological process (BP) branch of the gene ontology (GO) database revealed a significant downregulation of genes associated to extracellular matrix (ECM) organization and cellular adhesion, while an upregulation of genes in the immune response category (adj. p-value < 0.01, **Fig. 4C)**. Interestingly, we observed the downregulation of ECM genes and niche factors, such as *TnC*, *Pf4*, and *Cxcl12* **(Fig. 4D)**, thus suggesting an impaired supportive activity for HSC. Additional genes, such as *Sparc*, *Col1a2*, *Tgfb1*, *Fgf1* and *Kitl*^25^ showed a trend of downregulation **(Suppl. Fig. 8)** in *th3* Mk.

**Figure 4.**
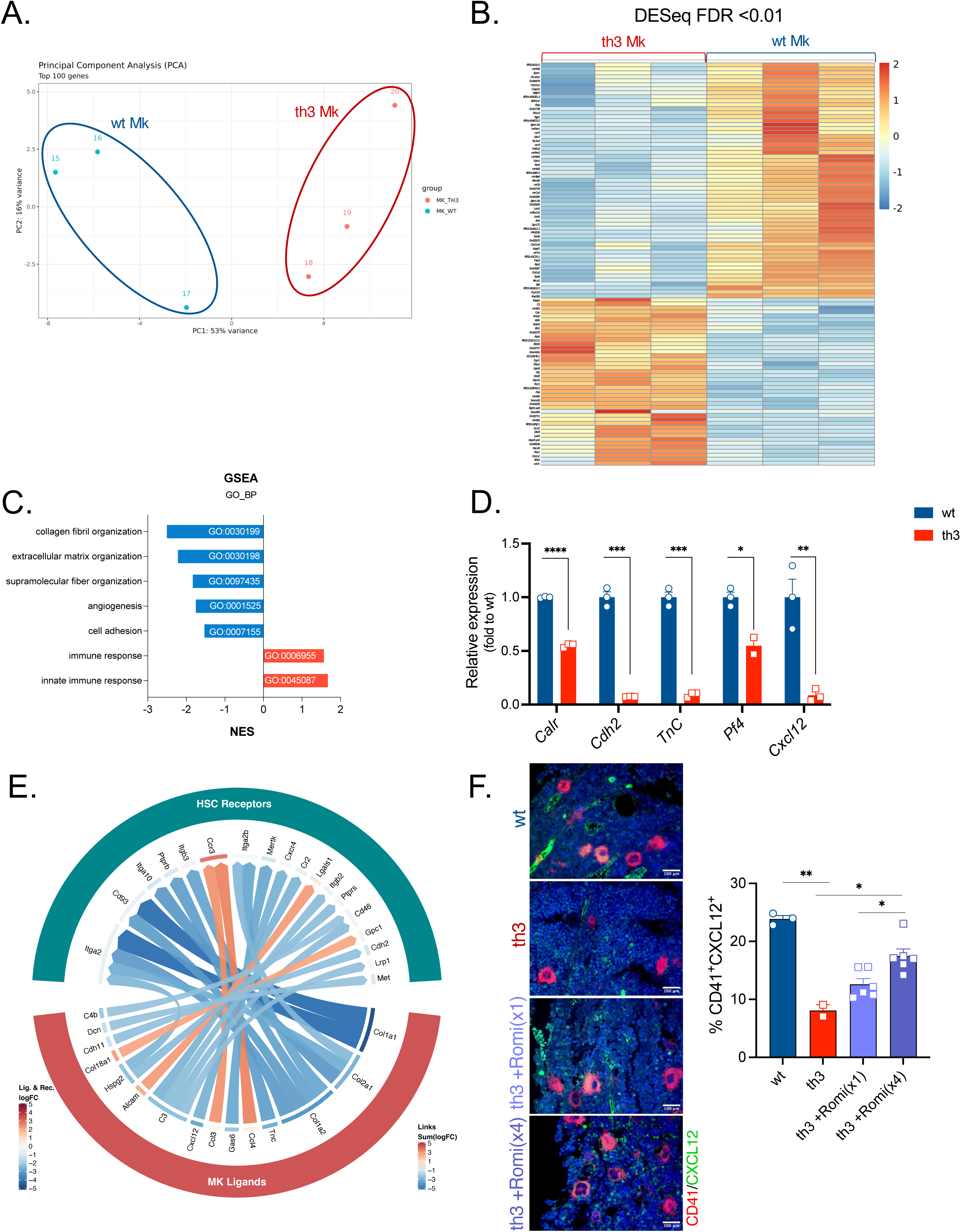
Altered gene expression profile and defective BM niche support by BThal Mk. **A.** Principal Component Analysis (PCA) using the top 100 most variable genes in *th3* compared to wt sorted Mk by RNA-seq (wt *n* = 3; *th3 n* = 3). **B.** Heatmap of DEG (FDR<0.01) in wt and *th3* Mk samples (wt *n* = 3; *th3 n* = 3). **C.** Gene set enrichment analysis (GSEA) employing the biological process (BP) branch of the gene ontology (GO) database. Normalized enrichment scores (NES) of selected categories are reported. **D.** Expression levels of extracellular matrix and niche factors in wt and *th3* sorted Mk by RNA-seq analysis. Data are reported as normalized counts and expressed as fold to wt controls (wt *n* = 3; *th3 n* = 3). **E.** Chord diagram showing the top 25 ligand-receptor pairs obtained from the bioinformatic analysis of the RNA-seq data on sorted Mk (ligands) and HSC (receptors) samples, from wt and *th3* mice. Each line connecting ligand-receptor pairs represents an overexpressed (red) or underexpressed (blue) interaction in *th3* vs. wt populations, according to the combined score (computed on logFC values). Logaritmic fold change (LogFC) expression of ligand and receptor is reported close to gene names, according to the color scale bar. **F.** Histological analysis (immunofluorescence) to confirm Mk-HSC interactions (wt *n* = 3; *th3 n* = 2; *th3*+Romi(x1) *n* = 6; *th3*+Romi(x4) *n* = 6).

Given the direct role that Mk play on HSC function within the BM niche, we exploited a bioinformatic analysis of the interactome^33^ to assess whether the HSC-Mk niche crosstalk was altered. We scored the ligand-receptor pairs present in a published database by using a combined score computed on the logFC values of the comparisons between *th3* and wt samples, in both Mk and HSC. Among the top 25 selected pairs, we found downregulated key niche interactions, such as *Cxcl12-Cxcr4*, *TnC*-*Prprb* and *Cdh11*-*Cdh2*, thus suggesting an altered HSC-Mk crosstalk **(Fig. 4E and Suppl. Table 2)**. Immunofluorescence analysis of CXCL12 protein expression by CD41^+^ Mk confirmed the transcriptomic data of reduced levels in *th3* Mk that were restored upon Romiplostim treatment **(Fig. 4F)**.

## DISCUSSION

In the last decade the use of transgenic models knocking-out or overexpressing key HSC or niche molecules helped to dissect the complexity of the HSC-BM niche crosstalk, in steady state or upon acute stimulation^3^. However, HSC-niche interactions are still underexplored in disease conditions associated to chronic stress. BThal is characterized by a chronically altered BM microenvironment with changes in the levels of multiple local and systemic factors including reduction of systemic TPO^10,11,13,15^. We exploited this model to define the role of TPO signaling on HSC and the BM niche, overcoming limitations of using constitutive and conditional TPO or MPL knock-out experimental systems. TPO indeed is a key regulator of both HSC function^18,19^ and Mk differentiation leading to Plt production^17,34^. Mk have been described as active components of the HSC niche^6,25,35^ but no studies so far elucidated the role of TPO on the BM niche. Our work highlighted how reduced TPO caused by defective Plt clearance, affects HSC function and the megakaryocytic BM niche **(Fig. 5)**.

**Figure 5.**
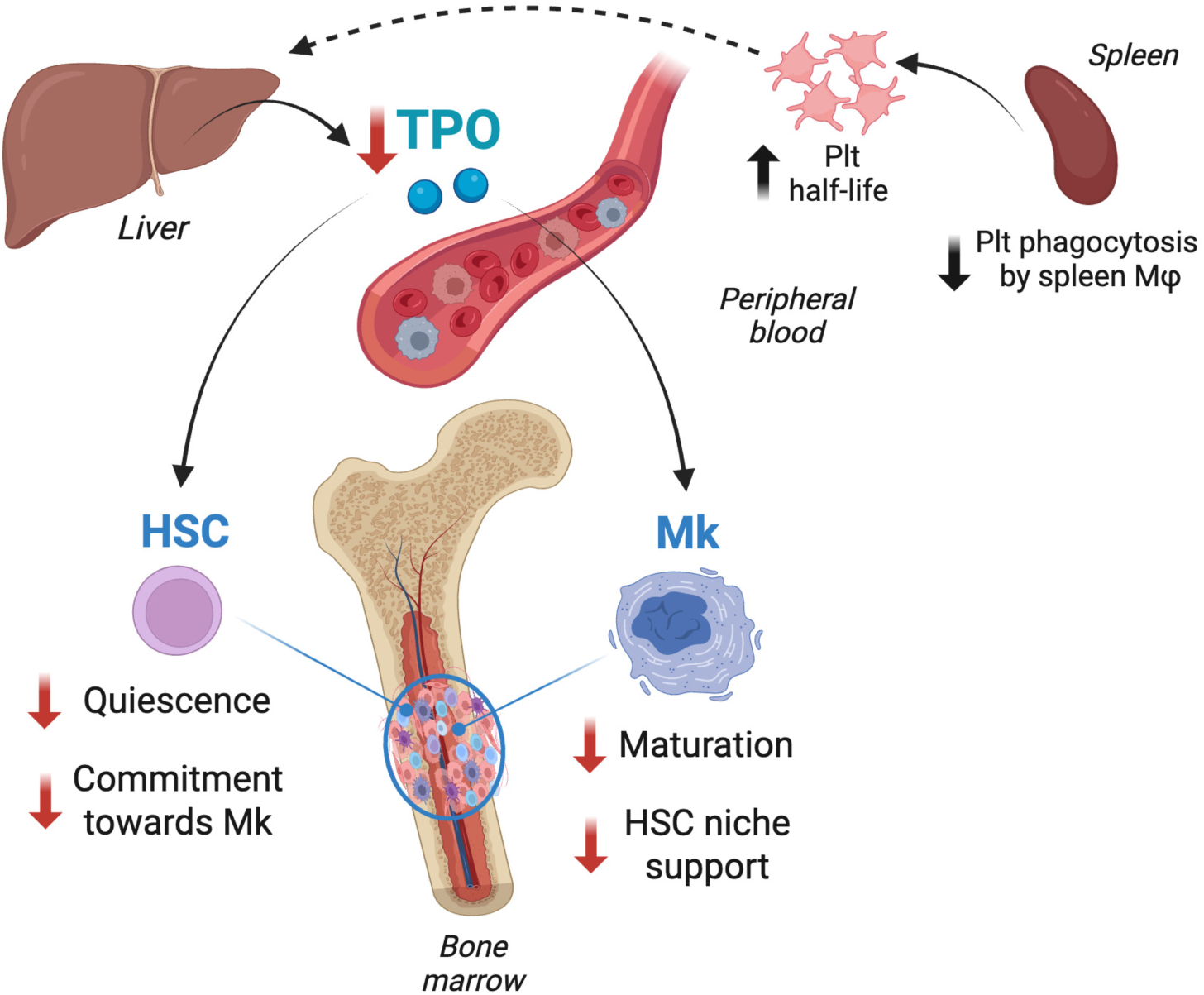
Working model of the dual effect of chronically reduced TPO on HSC and Mk BM niche in the BThal model. In *th3* mice the reduced clearance of Plt by splenic Mϕ increases Plt half-life in circulation, causing the decreased production of TPO by liver. Chronically low TPO has a dual effect on HSC, reducing their quiescence and commitment towards Mk, and on Mk, by impairing their maturation and HSC niche supporting capacity.

First, we investigated the origin of reduced TPO in Bthal *th3* mice and we demonstrated that is caused by the prolonged half-life of circulating Plt which accumulate in the circulation because of a reduced clearance by spleen Mϕ. Since iron massively accumulates in BThal spleen^36^ and in particular in Mϕ, we believe that excessive iron causes an impairment of their phagocytic activity, as described in models of iron overload^37^.

Low TPO levels in *th3* mice causes a reduced response to TPO in HSC resulting in downregulation of target genes associated to stemness, including *Cdkn1a*, *Hoxa9* and *Hoxb4*^18,19,38^. Moreover, TPO defect impacts on the commitment of HSC towards the Mk lineage, with under-expression of Mk-biased genes and lower frequency of CD41^+^ and CD9^high^ HSC. TPO indeed was reported to be responsible for maintenance of HSC with megakaryocytic differentiation potential^39,40^. Notably, *in vivo* stimulation of TPO axis in *th3* mice restored the pool of quiescent HSC, thus proving the contribution of defective TPO signaling in altering HSC.

Since TPO is a key regulator also of Mk maturation^34^, we investigated the effect of chronically reduced TPO on megakaryopoiesis and we observed a defect in Mk maturation both in *th3* mice and BThal patients, with loss of the mature polyploid profile. These findings correlated with a reduced *in vivo* Plt biogenesis and impaired *in vitro* differentiation of Mk. To reconcile these results with the higher Plt count we showed that, despite the reduced generation of Plt, they accumulate in the circulation because of their prolonged half-life and reduced clearance.

The maturation defect of *th3* Mk results in a reduced expression of ECM and niche factors, suggesting a reduced function in supporting BM niche and HSC function. ECM not only provides structural support in the BM, but it is also a critical regulator of HSC behaviour. Mk produce ECM components with an active role in regulating megakaryopoiesis and the BM microenvironment^28,41–43^. We found significantly reduced expression levels of *Tnc* encoding for the glycoprotein tenascin C which is known to affect HSC migration, adhesion and reconstitution potential^44^. Reduction of *Pf4* and *Cxcl12* in *th3* Mk are consistent with an impaired supporting activity to HSC. PF4 was reported to be a key regulator of the megakaryocytic niche, important for the maintenance of HSC quiescence^6^. These data reflect what already observed in other conditions associated to dysmegakayopoiesis affecting HSC maintenance^26^. We did not consider local TPO production in the BM by Mk as a potential feedback loop, given the consensus knowledge that systemic TPO secreted by hepatocytes is the one required for HSC maintenance^24^. Interestingly, the positive enrichment in immune response categories of the genes expressed by *th3* Mk might be associated to an immunomodulatory role in this disease context or to the expansion of the immune Mk subset in presence of reduced TPO levels^27,28^.

Interactome analysis of RNA-seq data from sorted *th3* Mk and HSC revealed an altered crosstalk, with downregulation of key ligand-receptor pairs, such as *Cxcl12-Cxcr4* at transcript and protein levels. Our results are consistent with the recent identification by transcriptomic and functional assays of a niche-supporting subpopulation within the Mk heterogeneity. Niche-supporting Mk were demonstrated to be mature high ploidy Mk^27^ and our data showed that the chronically reduced TPO stimulation actually causes a reduced maturation of Mk. Furthermore, in our model the TPO defect generates a lower proportion of Mk-biased HSC and niche Mk are derived from Mk-biased HSC through a direct differentiation route^28^. Overall, our data are in line with the most recent findings about the origin of niche-supporting Mk obtained in other experimental settings at steady-state condition.

We propose that the impact of TPO on HSC is both direct, since HSC express MPL receptor, and indirect *via* the megakaryocytic BM niche. In this view, the seminal papers about the role of TPO on HSC based on TPO or MPL constitutive and hepatic TPO knock-out models^18,19,24^ should be considered under a new perspective, that is not only the direct effect of TPO on HSC, but also the indirect one through the niche-supporting Mk. Moreover, very recently TPO has been identified as a pivotal systemic regulator of the total number of HSC in the body^20^, although the role of the Mk niche in this mechanism has not been assessed. Further studies will elucidate the extent of the contribution of the direct and indirect TPO effects on HSC number and function.

We provided a model of genetic anemia as an ideal setting to study TPO regulation, overcoming the experimental limitations of the transgenic models used so far. If on one hand the *th3* mouse represents a “real-world” disease setting of altered BM niche and HSC function^10,11,13^, on the other side the implications of our findings on BThal pathophysiology require further investigation. Plt accumulation in BThal patients is reported to be associated to Plt hyperactivated state and hypercoagulability^29,30^ and we showed here reduced Mk maturation in patient-derived BM biopsies, however the TPO defect has not been reported in BThal. In the multifactorial secondary complications associated to the disease the role of altered megakaryopoiesis might be caused by different factors. The iron overload condition associated to the disease, for example, might contribute to the defect since low iron was shown to promote Mk differentiation^45^. Nevertheless, our findings might be relevant for better understanding diseases associated to altered TPO regulation or treatment with TPO-ra. Indeed, patients with loss-of-function mutations in MPL or TPO develop congenital amegakaryocytic thrombocytopenia and subsequent bone marrow failure^22,46^.

Overall, our study reconsidered the role of TPO on hematopoiesis providing three main aspects of novelty: 1) we unraveled the role of TPO on the megakaryocytic BM niche; 2) we highlighted the dual effect of TPO through a direct mechanism on HSC and an indirect one mediated by the niche-supporting Mk; 3) we exploited a natural model of genetic anemia as ideal condition of chronically low TPO stimulation, caused by the reduced clearance and prolong half-life of circulating Plt.

We consider as limitation of this work some missing evidence in humans, since most of the study was performed in the mouse model. Indeed, the collection of adequate numbers of BM bioptic samples from patients and normal subjects has intrinsic limitations since it is not ethically justified for research use. Moreover, we cannot exclude any indirect role of TPO on other niche populations, such as bone cells. Megakaryopoiesis indeed has been demonstrated to be part of a complex cross-regulation with the BM endosteal niche, as Mk control also bone homeostasis^47–49^.

Altogether these data showed that TPO affects both HSC and the BM niche in a model of chronically reduced stimulation, paving the way towards a more comprehensive knowledge of direct and indirect mechanism driven by systemic TPO in the BM microenvironment.

## METHODS

### Mice

Male and female C57BL/6 and C57BL/6-CD45.1 (B/6.SJLCD45a-Pep3b) wild-type (wt) mice were purchased from Charles River. C57BL6/Hbb^th^^3/+^ mice lack both the b1 and b2 adult globin genes^16^, resulting in perinatal death in homozygosity but showing characteristics of severe BThal intermedia in heterozygosity. C57BL6/Hbb^th^^3/+^ mice were purchased from The Jackson Laboratory and bred in the laboratories at San Raffaele Hospital to maintain the colony in heterozygosity. All the analyses were performed on adult 12-14 weeks old mice. Experimental and control mice were age and sex matched littermates for individual experiments. All animal experiments were performed in accordance with approved protocols of the Institutional Animal Care and Use Committees of San Raffaele Institute (IACUC #735 and 1047).

### Human samples

BM sampling was performed for pre-transplant marrow evaluation in donors and patient candidates to allogeneic bone marrow transplantation^50^. For control purposes, BM bioptic samples from patients with secondary polycythemia were selected. All human samples were obtained after informed consent from patients or legal guardians and with the approval of Institutional Ethical Committees

### In vivo experiments

For acute Plt depletion and *in vivo* Plt biogenesis experiments, Plt depletion was performed by intravenous injection of 0.5 mg/g of rat purified anti-mouse GPIba (Emfret Analytics) in PBS, as previously reported^51^. At different time points after injection, blood has been collected from the retro-orbital sinus to analyze Plt counts by using a hemocytometer.

To evaluate *in vivo* Plt half-life, Plt clearance was evaluated by retro-orbital injection of 3 mg of DyLight 488–labeled anti–GPIb-V-IX (Emfret Analytics) antibody diluted in PBS, as as previously reported^51^. To analyze the percentage of labeled platelets, 10 mL of whole blood was incubated with 2 mL of anti-CD61 for 15 minutes, followed by addition of 150 mL of PBS. *th3* mice received subcutaneous injections one dose or 4 doses every 3 days of either vehicle (PBS) or Romiplostim (20 μg/kg of body weight; Amgen Inc) and were analyzed after 3 and 14 days, respectively. Doses and administration schedules were selected according to previous studies in mice^52,53^.

### Flow cytometry

Murine total BM cells were extracted from femurs and tibiae. Briefly, after cutting the epiphyses, the BM was flushed out by using a 1 ml syringe filled with ice cold MACS buffer supplemented with BSA and EDTA (Milteny Biotech) and the cell suspension was filtered through a 70 μm cell strainer. To analyze primitive progenitors and stem cell populations, Lineage^-^ cells were isolated from total BM samples as a negative selection by depletion of cells expressing a panel of lineage markers by mouse Lineage Cell Depletion Kit (Milteny Biotech), following manufacturer instructions. The unlabeled Lineage^-^ cells were collected. All staining was performed with commercially prepared antibodies in MACS Buffer.

Flow cytometry analysis was performed with freshly isolated BM cells. For HSC, MkP, PreMegE analyses, Lineage^-^ cells, enriched in stem and progenitors, were stained with Lin-Alexa700 (BioLegend), CD117-APC (BD Pharmingen) or CD117_APC-Cy7 (Biolegend), Sca1-PE-Cy7 (BD Pharmingen), CD48-PB (BioLegend), CD150-PE (BD Pharmingen) or CD150-BV421 (BD Horizon) or CD150-APC (BioLegend), CD41-FITC (BD Pharmingen), CD9-BV510 (BD Optibuild), CD16/32-PerCP-Cy5.5 (BD Pharmingen), CD105-PE (BioLegend) and Fc Block (BD Pharmingen). Mk staining was performed on freshly isolated BM cells by Lin-PB (BioLegend), CD41-FITC (BD Pharmingen) and CD61-PE (BD Pharmingen). Mk absolute counts were performed by adding CountBrightTM absolute counting beads (Invitrogen), according to manufacturer’s protocol.

All the surface staining were performed with commercially prepared antibodies in MACS Buffer at 4°C for 30 min. Cell cycle analysis of HSC was performed by staining Lin^-^ cells for surface markers and 7-AAD after fixation and permeabilization protocol (BD Pharmingen). *Ex vivo* Mk ploidy was analyzed by Draq5 (Biostatus) staining. For *in vitro* Mk ploidy, cells were fixed overnight in ice cold 70% ethanol at -20°C. Sample were incubated in PBS with 100 μg/ml of RNAse and propidium iodide solution and stained with FITC anti mouse CD41 (MWReg30, Biolegend). MkP and PreMegE were stained as previously reported^32^. Spleen cells were stained by F4/80-APC (eBioscience) and with 0.125 μM calcein-acetoxymethyl ester (Calcein-AM, Invitrogen) in PBS for 5 min at 37°C. Then cells were washed with DPBS and stained for F4/80 surface markers. Finally, calcein-AM fluorescence was analysed by flow cytometry. Spleen Mϕ were stained using F4/80-PE (eBioscience), CD11b-FITC (BD Pharmingen), VCAM1-BV711(BD Optibuild) antibodies.

All antibodies were titrated and validated using appropriate positive and negative control samples. Unstained cells, single stained compensation beads (BD Biosciences) and fluorescence minus one (FMO) were used for compensation and as a control to set the appropriate gates, respectively. Flow cytometry analyses were performed on FACS Canto II (BD Biosciences) and analyzed with FCS Express Software (De Novo Software).

### Platelet collection and analysis

Blood collected in Citrate Phosphate Dextrose Solution (Sigma) from the retro-orbital sinus was analyzed with a Sysmex KX-21N hemocytometer (Sysmex). To collect platelet-rich plasma, blood was then diluted with the same volume of 13 Tyrode buffer (5 mmol/L HEPES, 137 mmol/L NaCl, 2.7 mmol/L KCl, 0.4 mmol/L NaH2PO4, and 2.8 mmol/L dextrose, pH 7.4) and centrifuged for 7 minutes at 700 rpm, as previously reported^51^.

For Annexin V staining, 200,000 Plt were stained in 100 mL of 13 Binding Buffer with 5 mL of phycoerythrin–Annexin V (BD Pharmingen) for 15 minutes at room temperature, followed by stopping reaction by 100 mL of Binding Buffer addition. To analyze CD62P and CD47 surface expression, 5 mL of PRP were stained with anti-mouse CD61 and anti-mouse CD62P or CD47. To analyze CD62P surface expression, 5 mL of PRP was incubated with thrombin (final concentration, 1 U/mL; Sigma-Aldrich) for 2 minutes at room temperature.

### In vitro Plt phagocytosis

We performed *in vitro* phagocytosis assay by incubating wt or *th3* Plt labelled with CellTrace FarRed^TM^ with total splenocytes from wt or *th3* animals. Matched (wt Plt + wt splenocytes; *th3* Plt + *th3* splenocytes) and mismatched (wt Plt + *th3* splenocytes; *th3* Plt + wt splenocytes) settings were evaluated and the frequency of splenic Mϕ that had phagocytosed Plt was assessed by analyzing CellTrace FarRed^+^ F4/80^+^ CD11b^+^ VCAM^+^ cells by flow cytometry.

### In vitro Mk differentiation

For differentiation of mouse Mk, BM cells were incubated with FITC Lineage cocktail antibodies and anti-FITC microbeads-conjugated antibodies according to the manufacturer’s instructions (Miltenyi Biotech). Lineage^-^ fractions were cultured for 4 d in DMEM (Gibco) supplemented with 1% of penicillin/streptomycin, 1% of L-glutamine, and 10% of fetal bovine serum (Gibco), in the presence of 10 ng/ml of murine recombinant TPO (Peprotech), as previously reported^52^.

CFU-Mk assay was performed using MegaCult^TM^ medium (Stemcell Technologies), a collagen-base medium for colony assay of Mk progenitors, following manufacturer instructions.

### Histology and immunolocalization analyses

Bone samples were dissected from mice, washed in PBS and collected for fixation in 10% neutral buffered formalin and embedded in paraffin. Decalcification was performed using an EDTA-based decalcifying solution (Kaltek). In detail, samples were decalcified for 8 hours, washed in water for 1 hour, and subsequently processed for paraffin embedding.

Four-micrometers-thick human and mouse tissue sections were stained with H&E for histological evaluation. For double-marker immunofluorescence (IF), tissue sections were deparaffinized, rehydrated, and unmasked using Novocastra Epitope Retrieval Solutions at pH9 (Leica Biosystems) in a thermostatic bath at 98°C for 30 minutes. After cooling to room temperature and washing in PBS, nonspecific Fc-mediated binding was blocked with 0.4% casein in PBS (Leica Biosystems). Sections were then incubated with CD41 (clone EPR17876, 1:250 pH9, code ab181582, Abcam) and CXCL12 (clone SDF1, 1:50 pH9, code 79018, R&D) primary antibodies. The binding of the primary antibodies to their respective antigenic substrates was revealed by made-specific secondary antibodies conjugated with Alexa-488 (Life Technologies, 1:250) and Alexa-568 (Life Technologies, 1:300) fluorochromes. The slides were analyzed under a Zeiss Axioscope A1 microscope equipped with four fluorescence channels widefield IF. Microphotographs were collected using a Zeiss Axiocam 503 Color with the Zen 2.0 Software (Zeiss).

Quantitative analyses of Mk diameters and IF stainings were performed using HALO image analysis software (v3.2.1851.229, Indica Labs). CD41/CXCL12 double-positive cells were quantified by calculating the percentage of CD41^+^/CXCL12^+^ cells in five non-overlapping high-power fields (x400).

### ELISA

ELISA for murine TPO (Quantikine®, R&D Systems) was assayed on sera from wt and *th3* mice, according to the manufacturer’s instructions. *ddPCR gene expression*

Total RNA was isolated from hepatocytes by AllPrep DNA/RNA Micro Kit (QIAGEN). RNA was reverse-transcribed using Precision nanoScript 2 RT kit (Primer Design). Runs were performed on QX200™ Droplet Digital™ PCR (ddPCR™) System (Biorad), by using ddPCR™ Supermix for Probes (No dUTP) and PrimePCR™ ddPCR™ Expression Probe Assay, following the manufacturer’s protocol. PrimePCR™ ddPCR™ Expression Probe Assay (Biorad): Tpo (qMmuCEP0039012), b-actin (qMmuCEP0039589).

### RNAseq analysis

Total RNA from HSC sorted from pool of mice and Mk sorted from single animals was isolated by AllPrep DNA/RNA Micro Kit (QIAGEN). Library preparation and RNA-seq analysis was performed as described in^10^. Briefly, Ovation SoLo RNA-seq kit (Illumina) has been used for library preparation following the manufacturer’s instructions. Libraries were processed with Illumina cBot for cluster generation on the flowcell and sequenced on paired-end 125 bp mode on HiSeq2500 (Illumina). The quality of the RNA-seq data was determined using FastQC and low-quality sequences were trimmed using trimmomatic. Reads were aligned to the *Mus musculus* reference genome (GRCm38/mm10) and filtered considering the Unique Molecular Identifiers (UMI) tags. Transcript UMI counts were normalized for library size using DESeq2’s median of ratios in R using the DESeq2 package, and differential gene expression analysis between wt and *th3* mice was performed considering *p-values* corrected by FDR. To determine whether some specific gene sets were significantly different between wt and *th3* mice, we performed Gene Set Enrichment Analysis (GSEA) employing the biological process (BP) branch of the gene ontology (GO) database on genes ranked by log2 fold change values. Relative expression data of selected genes were reported as gene UMI counts relative to *Hprt* and shown as fold to wt controls.

Interactome analysis was performed exploiting a publicly available database of ligand-receptor pairs^33^. In details, for each pair, the logFC values resulting from the DGE analysis between *th3* and *wt* mice in Mk samples were used to score ligands, while those obtained from the HSC ones were used for receptors. The final score of each pair was then obtained by summing the two corresponding logFC values, and ligand-receptor pairs were ranked according to the absolute score value.

### Statistics

No statistical methods were used to predetermine the sample size. Sample sizes were determined based on previous studies from our laboratory^10,11^. Analyses were performed by using GraphPad prism v8.0 software (La Jolla, CA, USA). All data are presented as single values and mean±SEM. Mann-Whitney test was used for comparisons between two independent groups, Bonferroni correction was applied for comparison among more than 2 groups. Statistical significance is reported as follows: * *p* < 0.05, ** *p* < 0.01, *** *p* < 0.001, **** *p* < 0.0001.

### Data availability

Gene expression RNA-seq data for this study have been deposited in the European Nucleotide Archive database with accession numbers PRJEB31882 for HSC samples (http://www.ebi.ac.uk/ena/data/view/PRJEB31882) and PRJEB98475 for Mk samples (http://www.ebi.ac.uk/ena/data/view/PRJEB98475).

## AUTHOR CONTRIBUTIONS

A.A. designed the experimental work, performed the research, analyzed, interpreted and discussed data, provided project coordination, obtained financial support, and wrote the paper; A.M. provided key reagents, performed *in vitro* Mk differentiation, contributed to design the experimental work, discussed data and critically reviewed the paper; M.S., S.S. and L.R. contributed to experimental work; A.G., V.C., G.M., C.T. and M.P. performed histological analysis; S.B. and I.M. performed bioinformatic analysis on the RNA-seq data; S.M. provided human samples; C.T. and A.B. discussed data and critically reviewed the paper; G.F. supervised the project, obtained financial support, discussed data, and wrote and critically reviewed the paper; and all authors approved the final version of the manuscript.

## FUNDING SUPPORT

This work was supported by Telethon SR-TIGET Core Grant 2021 (G.F and A.A.), the European Hematology Association Junior Research Grant 2019 (A.A.) and the Italian Association for Cancer Research (AIRC) Investigator Grant 28978 (C.T.). A.A. is supported by the Cooley’s Anemia Foundation Research Fellowships 2018 and 2019, the American Society of Hematology (ASH) Global Research Award 2021 and by the Cariplo Foundation Young Researchers Grant (2022-0409).

## AKNOWLEDGMENTS

The authors thank the Flow Cytometry Resource, Advanced Cytometry Technical Applications Laboratory (FRACTAL) at Ospedale San Raffaele for cell sorting and the University Centre of Statistics in the Biomedical Sciences (CUSSB) at Vita-Salute San Raffaele University for help with the statistical analyses. We thank Amgen Inc for providing Romiplostim-Nplate®.

## CONFLICT-OF-INTEREST STATEMENT

The authors have declared that no conflict of interest exist.

## Notes

### Competing Interest Statement

The authors have declared no competing interest.

