## Supplementary Material for "A genetic anemia disentangles the dual effect of chronically reduced thrombopoietin on hematopoietic stem cells and bone marrow niche"

### Supplemental Tables

**Supplementary Table 1: DEG of *th3* Mk compared to wt ones.** Differentially expressed genes (DEG) by DESeq analysis of RNA-seq data in Mk from *th3* vs. wt mice

| Gene | logFC | PValue | FDR |
|---|---|---|---|
| RP23-8J15.5 | 4,68296424 | 3,314E-107 | 4,482E-103 |
| mt-Nd3 | -4,7295725 | 1,1666E-84 | 7,8891E-81 |
| Sparc | -3,2246518 | 1,5614E-16 | 7,0393E-13 |
| Clca3a1 | 1,81910027 | 5,0356E-16 | 1,7027E-12 |
| Gm26870 | -2,2605724 | 2,3631E-15 | 6,3921E-12 |
| Col11a1 | -4,1660918 | 1,1746E-11 | 2,6476E-08 |
| Casp12 | 1,83060992 | 2,3977E-11 | 4,6327E-08 |
| Adgrl4 | 1,5835105 | 2,8439E-11 | 4,808E-08 |
| Aldh1a1 | 1,68064017 | 3,5277E-10 | 5,3014E-07 |
| Gm21738 | -2,2908417 | 1,2059E-09 | 1,631E-06 |
| Plscr2 | 3,19441759 | 1,41E-09 | 1,7337E-06 |
| Iigp1 | 2,35793942 | 2,2036E-09 | 2,4836E-06 |
| Ibsp | -3,3153364 | 5,3312E-09 | 5,5465E-06 |
| mt-Rnr1 | -1,1714202 | 6,4531E-09 | 5,8186E-06 |
| mt-Ti | -1,9988333 | 6,0241E-09 | 5,8186E-06 |
| RP24-459O21.3 | -1,3433618 | 8,9816E-09 | 7,5922E-06 |
| Uba7 | 0,87781656 | 2,2018E-08 | 1,7517E-05 |
| Slc2a3 | 0,87407323 | 3,399E-08 | 2,554E-05 |
| RP24-459E13.2 | -1,4893927 | 6,4326E-08 | 4,3501E-05 |
| Ighv1-34 | 21,1533285 | 6,2669E-08 | 4,3501E-05 |
| mt-Tt | -1,6300169 | 7,8767E-08 | 5,073E-05 |
| Col5a2 | -3,2386249 | 1,8531E-07 | 0,00011393 |
| mt-Rnr2 | -1,0610924 | 2,022E-07 | 0,00011395 |
| mt-Nd4l | -1,3487485 | 1,9451E-07 | 0,00011395 |
| Gvin1 | 3,06122264 | 3,2656E-07 | 0,00016988 |
| Rorb | 3,11327452 | 3,2577E-07 | 0,00016988 |
| mt-Nd2 | -0,9562165 | 3,4499E-07 | 0,00017282 |
| RP23-58B7.2 | -1,2880009 | 4,0593E-07 | 0,00019487 |
| mt-Atp8 | -1,3835514 | 4,1784E-07 | 0,00019487 |
| Rbm20 | -2,9349461 | 4,3639E-07 | 0,00019674 |
| mt-Tp | -1,9846912 | 4,696E-07 | 0,00020488 |
| Gm10718 | -2,4441153 | 5,0474E-07 | 0,00021201 |
| mt-Co2 | -1,2905997 | 5,1729E-07 | 0,00021201 |
| Gm22265 | -1,6282827 | 7,0602E-07 | 0,00028085 |
| Ighv1-26 | 4,17712017 | 8,7628E-07 | 0,00033862 |
| Gm22583 | -2,0173566 | 9,8436E-07 | 0,00036982 |
| Lars2 | -1,07703 | 1,0934E-06 | 0,00039968 |
| n-R5s115 | -1,8243156 | 1,1666E-06 | 0,00041229 |
| Jam2 | 2,28810174 | 1,1889E-06 | 0,00041229 |
| Art4 | 1,70135295 | 1,2343E-06 | 0,00041735 |
| Gpr171 | 1,34640806 | 1,3963E-06 | 0,00046062 |
| RP23-454A11.1 | 3,32979685 | 1,5374E-06 | 0,00049509 |
| Mir6236 | -1,2926721 | 1,6469E-06 | 0,00051179 |
| Tdrd9 | 3,19030088 | 1,665E-06 | 0,00051179 |

| | | | |
|---|---|---|---|
| Gm22973 | -1,6772654 | 2,1064E-06 | 0,0006331 |
| Col11a2 | -2,2899642 | 3,2966E-06 | 0,00096926 |
| Inppl1 | 1,33324856 | 3,5026E-06 | 0,00100792 |
| mt-Tw | -1,4208515 | 3,5802E-06 | 0,00100879 |
| RP23-42C15.1 | -1,7051478 | 4,014E-06 | 0,00110794 |
| Tnip3 | 1,17052876 | 5,0741E-06 | 0,00137253 |
| Nrp2 | -2,4567225 | 6,0235E-06 | 0,00159742 |
| Gm23287 | -1,4618336 | 6,3588E-06 | 0,0016539 |
| Col1a2 | -3,06231 | 6,6728E-06 | 0,00167129 |
| Eps8 | 1,92277246 | 6,6089E-06 | 0,00167129 |
| Abca1 | 1,02401802 | 7,1146E-06 | 0,00174954 |
| Skil | 1,01114879 | 8,3514E-06 | 0,002017 |
| RP23-362A12.3 | 1,10615184 | 1,0157E-05 | 0,00240911 |
| Tcp11l2 | 1,12182846 | 1,0331E-05 | 0,00240911 |
| Anp32b | -0,721165 | 1,0565E-05 | 0,00242194 |
| Enpp5 | 1,52994642 | 1,1125E-05 | 0,00250773 |
| C3 | -2,2026843 | 1,1533E-05 | 0,00255704 |
| mt-Nd1 | -0,7747666 | 1,23E-05 | 0,00268329 |
| Calr | -0,6153291 | 1,4424E-05 | 0,00309657 |
| Nr4a2 | 1,34565961 | 1,6865E-05 | 0,00356415 |
| Vldlr | 0,94364121 | 1,7247E-05 | 0,00358868 |
| Gatsl2 | 1,49896064 | 1,9736E-05 | 0,00404441 |
| Ifih1 | 0,73168914 | 2,0408E-05 | 0,00411967 |
| Gm22270 | -1,1769932 | 2,1383E-05 | 0,0041315 |
| Aspa | 1,46450597 | 2,1283E-05 | 0,0041315 |
| RP23-212C14.11 | -1,0072769 | 2,0887E-05 | 0,0041315 |
| Ifi203 | 0,7364003 | 2,2206E-05 | 0,00423008 |
| Gm10717 | -2,2915625 | 2,4312E-05 | 0,00456695 |
| Snord42a | -2,0019422 | 2,8151E-05 | 0,00521572 |
| AC124976.1 | -1,0544265 | 2,8795E-05 | 0,0052629 |
| Cyyr1 | 1,65247647 | 3,0872E-05 | 0,00556718 |
| Thbs1 | -1,0456683 | 3,4544E-05 | 0,00614747 |
| Irgm2 | 1,34841527 | 3,5282E-05 | 0,00619724 |
| Mt1 | 1,28237438 | 3,711E-05 | 0,00643476 |
| Oasl2 | 0,99092636 | 4,0312E-05 | 0,00681525 |
| Myo10 | -0,7805082 | 4,0002E-05 | 0,00681525 |
| Plcl1 | 2,11606033 | 4,1184E-05 | 0,00687668 |
| RP23-239P22.2 | -1,326353 | 4,2443E-05 | 0,00700047 |
| Flt4 | -2,5480491 | 4,4025E-05 | 0,00709376 |
| mt-Nd4 | -0,7676447 | 4,4057E-05 | 0,00709376 |
| Snora20 | -1,5963536 | 4,5039E-05 | 0,00716655 |
| Gm24245 | -1,0392737 | 4,6394E-05 | 0,0072962 |
| Rpl21-ps8 | -1,7992065 | 4,7676E-05 | 0,00741169 |
| Snord65 | -1,1677935 | 4,9871E-05 | 0,00766479 |
| Gm22711 | -1,4562361 | 5,3797E-05 | 0,00808457 |
| mt-Nd5 | -0,7859927 | 5,3332E-05 | 0,00808457 |
| RP23-52N2.1 | 1,42719512 | 5,5012E-05 | 0,00817627 |
| Isca1 | 0,73625937 | 5,9488E-05 | 0,00874539 |
| Dtx3l | 0,68362465 | 6,1849E-05 | 0,00899473 |
| Loxl2 | -4,0573743 | 6,431E-05 | 0,00925312 |

| | | | |
|---|---|---|---|
| Hspd1-ps3 | -1,1836852 | 6,5448E-05 | 0,00931774 |
| Gm25636 | -1,3704685 | 6,7083E-05 | 0,00945104 |
| Hacd4 | 1,01051418 | 6,8535E-05 | 0,0095561 |
| Ffar1 | -3,4675466 | 7,1601E-05 | 0,00966862 |
| Col1a1 | -4,630945 | 7,2202E-05 | 0,00966862 |
| Klhl4 | 2,50801731 | 7,1346E-05 | 0,00966862 |
| mt-Tc | -2,1520097 | 7,1808E-05 | 0,00966862 |

**Supplementary Table 2: Top 25 differential Mk-HSC interctions in *th3* and wt mice.** RNA-seq analysis of ligand expressed by Mk and receptors expressed by HSC in *th3* vs. wt mice

| Mk_Ligand | HSC_Receptor | Ligand_logFC | Receptor_logFC | Score |
|---|---|---|---|---|
| Col1a1 | Itga2 | -4,631182189 | -0,422648231 | -5,05383 |
| Col1a1 | Cd93 | -4,631182189 | -0,347297849 | -4,97848 |
| Col2a1 | Itga10 | -2,47249536 | -1,06600415 | -3,5385 |
| Col1a2 | Itga2 | -3,061661776 | -0,422648231 | -3,48431 |
| Col1a2 | Cd93 | -3,061661776 | -0,347297849 | -3,40896 |
| Tnc | Ptprb | -2,547009992 | -0,847188351 | -3,3942 |
| Col1a2 | Itgb3 | -3,061661776 | -0,155740237 | -3,2174 |
| Col1a2 | Itga2b | -3,061661776 | -0,061578543 | -3,12324 |
| Gas6 | Mertk | -2,040613612 | -1,04491708 | -3,08553 |
| Col2a1 | Itga2 | -2,47249536 | -0,422648231 | -2,89514 |
| Cxcl12 | Cxcr4 | -2,712059372 | -0,071022539 | -2,78308 |
| C3 | Cr2 | -2,1947949 | -0,497998349 | -2,69279 |
| Hspg2 | Itga2 | -2,177057599 | -0,422648231 | -2,59971 |
| Col2a1 | Itga2b | -2,47249536 | -0,061578543 | -2,53407 |
| C3 | Itgb2 | -2,1947949 | -0,240014995 | -2,43481 |
| Hspg2 | Ptprs | -2,177057599 | -0,245047806 | -2,42211 |
| C3 | Cd46 | -2,1947949 | -0,223306662 | -2,4181 |
| Cdh11 | Cdh2 | -1,18846277 | -1,100653223 | -2,28912 |
| C3 | Lrp1 | -2,1947949 | -0,079330065 | -2,27412 |
| Dcn | Met | -1,366954095 | -0,861187985 | -2,22814 |
| C4b | Cd46 | -1,997070062 | -0,223306662 | -2,22038 |
| Col18a1 | Gpc1 | 2,178792628 | 0,219195438 | 2,397988 |
| Alcam | Lgals1 | 1,988078918 | 0,665741735 | 2,653821 |
| Ccl3 | Ccr3 | 0,055644976 | 2,818788359 | 2,874433 |
| Ccl4 | Ccr3 | 0,393907868 | 2,818788359 | 3,212696 |

### Supplemental Figures

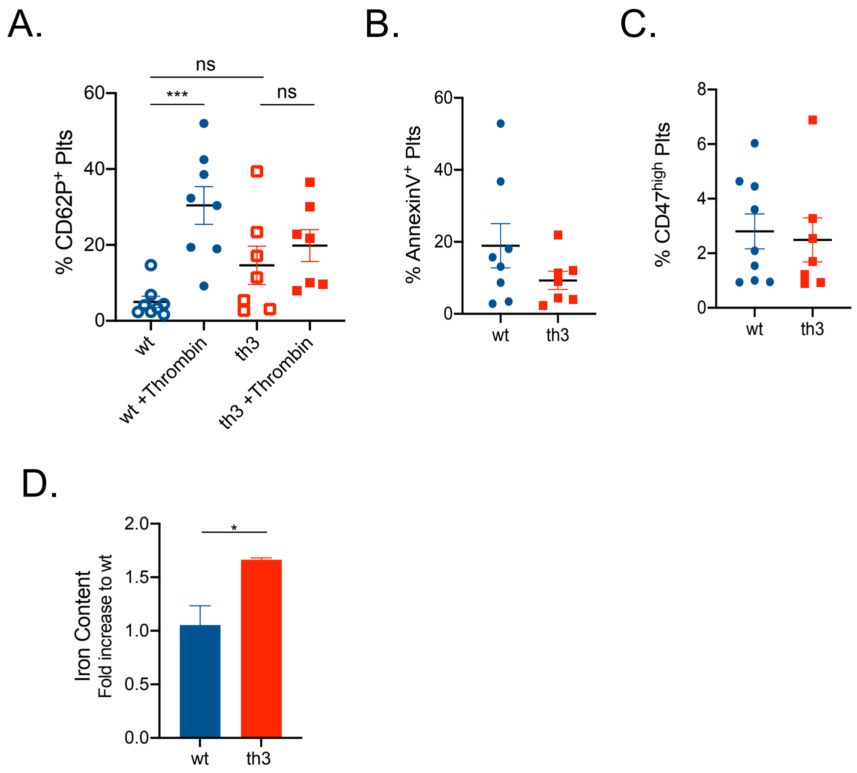


**Supplementary Figure 1. Plt surface markers and iron content of spleen Mφ in wt and *th3* mice. A.** Frenquency of $CD61^+$ Plt expressing CD62P, Annexin V and CD47 from wt and *th3* peripheral blood (wt $n > 6$; *th3* $n > 7$). **B.** Intracellular iron content in $F4/80^+$ Mφ from wt and *th3* spleens by flow cytometry analysis of calcein staining (wt $n = 3$; *th3* $n = 3$).

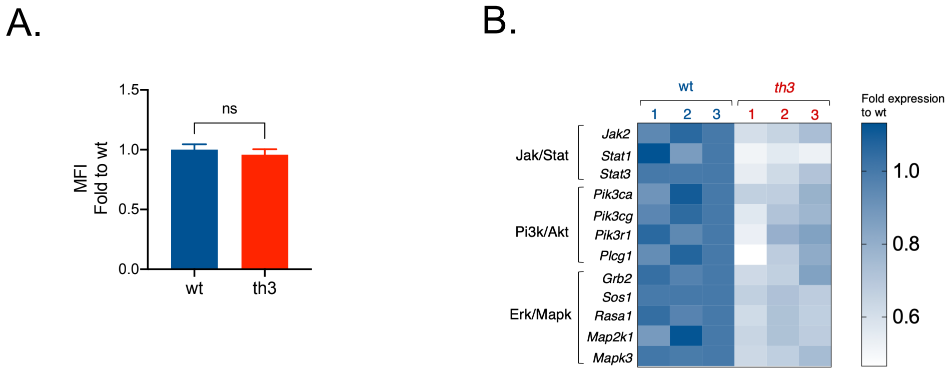


**Supplementary Figure 2. TPO signaling in wt and *th3* HSC. A.** Median fluourescence intensity (MFI) of surface MPL in wt and *th3* HSC by flow cytometry (wt *n* = 8; *th3 n* = 8). **B.** Gene expression heatmap of genes associated to TPO signaling in sorted wt and *th3* HSC by RNA-seq analysis. Data are reported as fold to wt controls (wt *n* = 3; *th3 n* = 3).

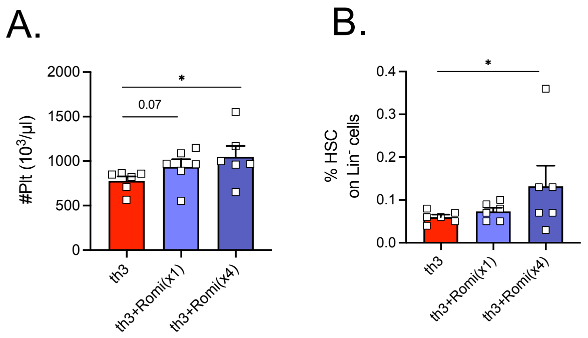


**Supplementary Figure 3. Plt count and HSC frequency upon *in vivo* stimulation of TPO axis by Romiplostim. A.** Circulating Plt count of *th3* mice treated with one or 4 doses of Romiplostim and controls by hemocytometer analysis (*th3* *n* = 6; *th3*+Romi(x1) *n* = 6; *th3*+Romi(x4) *n* = 6). **B.** Frequency of HSC on BM Lineage negative ($Lin^-$) cells of Romiplostim-treated and control *th3* mice (*th3* *n* = 6; *th3*+Romi(x1) *n* = 6; *th3*+Romi(x4) *n* = 6).

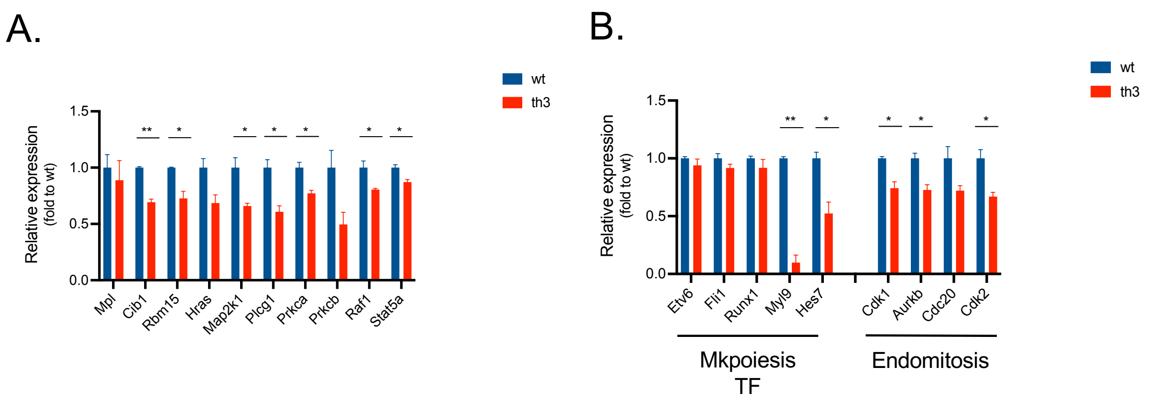


**Supplementary Figure 4. TPO signaling and megakariopoiesis genes in wt and *th3* Mk. A,B.** Expression levels of TPO signaling genes and genes involved in megakaryopoiesis and endomitosis in wt and *th3* Mk by RNA-seq analysis. Data are reported as normalized counts and expressed as fold to wt controls (wt $n = 3$; *th3* $n = 3$).

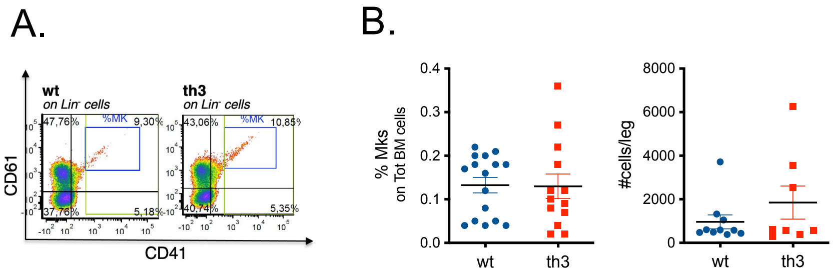


**Supplementary Figure 5. Analysis of BM Mk by flow cytometry analysis. A.** Gating strategy for the analysis of Lin$^-$ CD41$^+$ CD61$^+$ BM Mk. **B.** Frequency and absolute counts of Mk on total BM in wt and *th3* mice (wt *n* = 16 and 10; *th3 n* = 13 and 8, respectively).

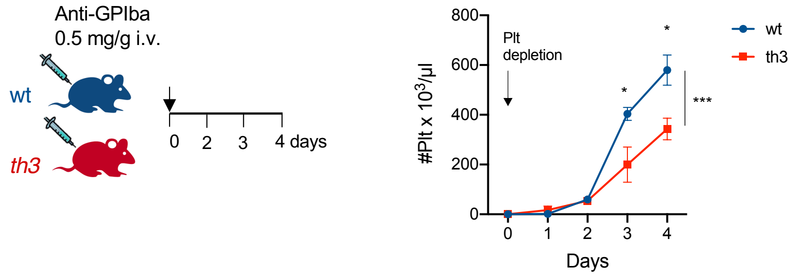


**Supplementary Figure 6. *In vivo* Plt biogenesis assay.** Plt depletion in wt and *th3* mice by anti-GpIba antibody. Plt recovery was assessed as Plt count at different time points after injection up to 4 days (wt $n = 4$; *th3* $n = 4$).

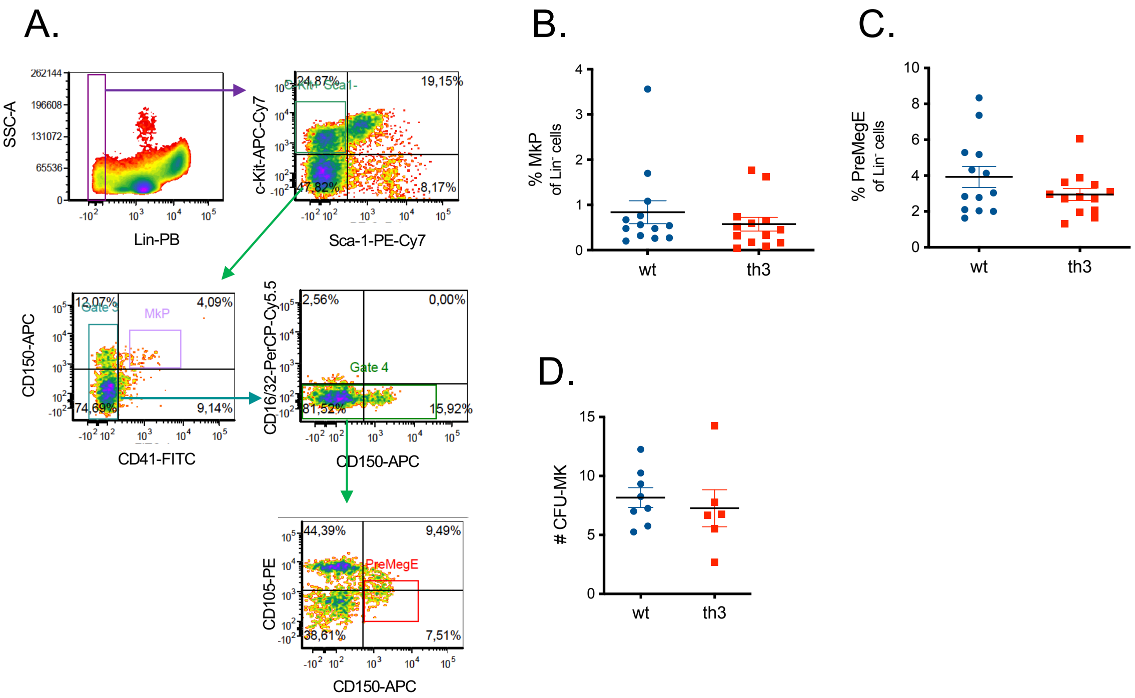


**Supplementary Figure 7. Analysis of BM Mk progenitors. A.** Gating strategy for the analysis of MkP and PreMegE BM populations. **B.** Frequency of MkP progenitors on Lin- cells in wt and *th3* BM by flow cytometry (wt *n* = 13; *th3 n* = 13). **C.** Frequency of PreMegE progenitors on Lin$^-$ cells in wt and *th3* BM by flow cytometry (wt *n* = 13; *th3 n* = 13). **D.** Number of CFU-Mk colony upon culture on methylcellulose of BM wt and *th3* cells (wt *n* = 8; *th3 n* = 6).

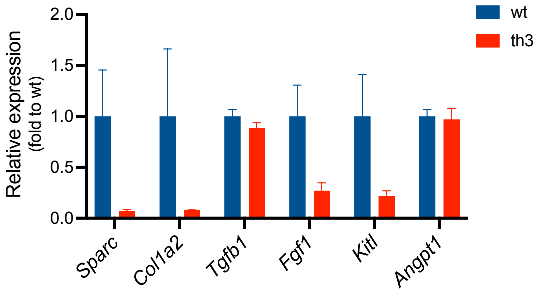


**Supplementary Figure 8. Altered expression of niche molecules by BThal Mk.** Relative expression levels of extracellular matrix and niche factors in wt and *th3* sorted Mk by RNA-seq analysis. Data are reported as normalized counts and expressed as fold to wt controls (wt *n* = 3; *th3* *n* = 3).
